# Library-Free Multiplexed Targeted Proteomics Enables Quantitative Protein-Level Genotyping

**DOI:** 10.1101/2025.09.10.675407

**Authors:** Steven R. Shuken, Steven P. Gygi

## Abstract

Existing targeted mass spectrometry methods require synthetic standard peptides, manual scan scheduling, or data libraries to quantify target proteins. Here we present GoDig-LiF, which uses spectra and retention times predicted by the Prosit-TMT model in place of data libraries, enabling targeted proteomics with only a tandem mass tag-labeled sample, target list, and mass spectrometer. We applied GoDig-LiF to the quantification of mutated proteins in cancer cell lines, including KRAS G13D.

## MAIN

Targeted mass spectrometry is one of the most powerful hypothesis-driven approaches to the identification and quantification of target proteins, having a higher sensitivity and dynamic range than untargeted MS.^1^ However, existing methods require synthetic standard peptides,^2,3^ manual scheduling of scans (as in parallel reaction monitoring [PRM]),^4,5^ or data libraries derived from deep proteomic analyses.^6,7,8^ GoDig is a targeted proteomic method that uses tandem mass tags (TMT) to increase sample throughput up to 35-fold, increase quantitative accuracy and precision, and increase data completeness relative to label-free methods. Although GoDig requires neither synthetic standards nor PRM scheduling, it requires a deep data-dependent acquisition (DDA)-based library; each time a new type of sample is to be analyzed, a new data library must be assembled, and library depth is limited by one’s ability to generate a large-scale sample, perform offline fractionation, and commit the required instrument time. Currently, there is no way to ensure that a new GoDig library includes a particular protein of interest.

To address this limitation, we used the Prosit-TMT model^9^ to assemble comprehensive sets of predicted spectra and elution orders (EOs) for GoDig analysis, resulting in an entirely DDA data-free workflow we call Library-Free GoDig (GoDig-LiF) (Fig. 1A). First, an *in silico* digest is performed with strict parameters, to manage the size of the predicted dataset while maximizing coverage; second, a more permissive *in silico* digest is performed on the target protein(s), ensuring that all possible precursor targets are included in prediction. Prediction is performed via the free-to-use Prosit website (https://www.proteomicsdb.org/prosit/); the GoDig-LiF software then converts the resulting outputs into the format of a GoDig library. This whole process takes less than one day with no sample preparation or instrument time required. There is complete flexibility with respect to species, organ, and cell type; any collection of protein sequences in FASTA format can be used.

**Figure 1.**
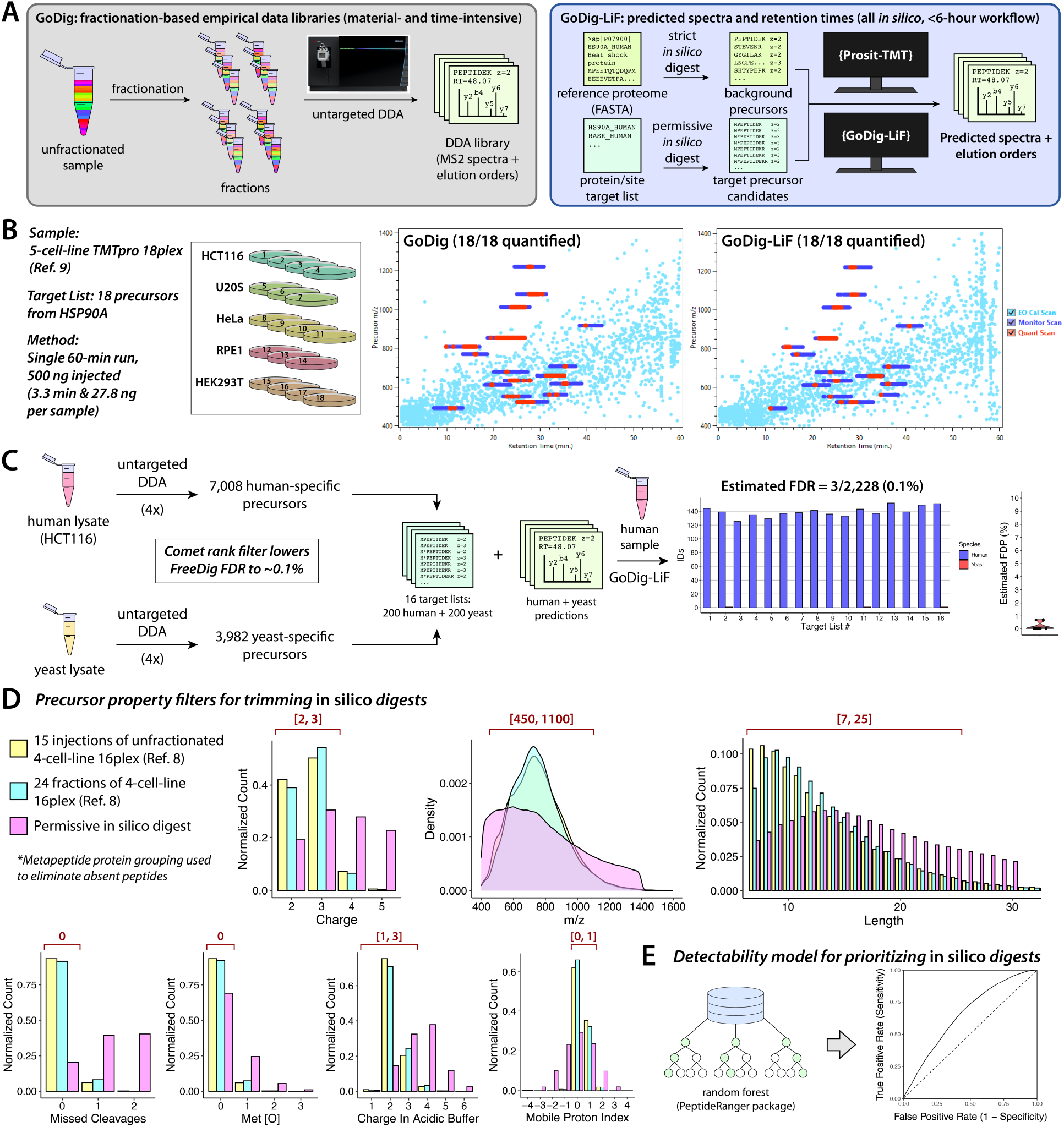
GoDig-LiF enables multiplexed targeted proteomics without synthetic standards, manual scheduling, or data libraries. **A.** Comparison between the GoDig library building workflow and the compilation of predicted spectra and elution orders in GoDig-LiF. **B**. Analysis of HSP90A peptides. **C**. Estimation of the identification false discovery rate (FDR) of GoDig-LiF. **D**. Comparison between properties of *in silico* predicted precursors and experimentally identified precursors, with efficient digestion filters indicated in red. **E**. Random forest-based model for TMT-labeled peptide detectability trained with the PeptideRanger R package.

To confirm the high accuracies of fragment ion intensity and retention time (RT) prediction by Prosit-TMT,^9^ we tested GoDig-LiF on a set of 18 peptides from the abundant chaperone protein HSP90A, using a previously described 4-cell-line 16plex sample (Fig. 1B, Table S1).^8^ The RT predictions were accurate enough to yield high-fidelity EO bins, yielding a 100% quantification success rate in a 60-min run. Manual inspection of spectra and EO bins confirmed prediction accuracy (Fig. S1–S2). To simulate a scenario in which a target protein’s detectable peptides are unknown, we performed a “semi-strict” *in silico* digest (see Methods) on HSP90A, resulting in 97 precursors, 62 of which are present in the largest publicly available GoDig library (Fig. S3, Table S2).^8^ With GoDig, we quantified 57 precursors corresponding to 30 peptides, and with GoDig-LiF, we quantified 58 precursors to 30 peptides, showing that even in a situation where excess undetectable precursors are targeted, GoDig-LiF can be performed without a loss in performance (Table S3).

To measure the false discovery rate (FDR) of GoDig-LiF, we used a two-species approach to identify peptides that were present and absent from the 4-cell-line mixture (Fig. 1C). By quadruplicate untargeted DDA analyses of a TMTpro-labeled yeast sample and human HCT116 cell sample, we created 16 target lists, each containing 400 targets (200 human-specific precursors and 200 yeast-specific precursors), ensuring that the precursors and peptide-spectrum matches had similar properties (Fig. S4, Table S4). Over 16 GoDig-LiF runs on the 4-cell-line 16plex, we initially measured a false discovery proportion (FDP) of 3.5 ± 1.1%, higher than the 1% FDR commonly accepted in untargeted proteomics (Fig. S5);^10^ to reduce this, we implemented a filter that requires the target to be the #1 ranking match to the spectrum according to the Comet search engine.^11^ As a result of this feature, the estimated FDR of GoDig-LiF is 0.1%, with an estimated 3 out of 2,228 identification events being false (Fig. 1C).

To reduce the number of precursors targeted per protein, we developed two tools: a set of filters based on precursor properties and a machine learning-based detectability model (Fig. 1D– E). To optimize the parameters of *in silico* digests, we compared a permissive human digest (see Methods) to two published datasets: a 24-fraction analysis of TMTpro-labeled peptides from four cell lines and a 15-injection analysis of an unfractionated 4-cell-line 16plex.^8^ We used the metapeptide protein grouping method^12,13^ to accurately identify peptides that should exist in the sample(s) (but are less detectable) based on the observed peptide set (Fig. S6). As a result, the strictest filters we recommend for target precursors are: length in [7, 25]; charge in [2, 3]; m/z in [450, 1100]; missed cleavages = 0; methionine oxidations = 0; charge in acidic buffer in [1, 2]; and mobile proton index in [0, 1], where the mobile proton index is defined as the charge minus the number of basic sites (H, K, R, and N terminus) in the peptide.^10,14^ To select detectable target precursors from filtered *in silico* digests, we used the PeptideRanger R package to train a random forest which estimates the detectability of a peptide from its sequence.^15^

We applied GoDig-LiF to a class of proteins that are absent from reference proteomes, and therefore from GoDig libraries, but present in cancer cell lines: mutated proteins (Fig. 2A). While DNA mutations are known to drive cancer, mutated proteins often play a role in producing the cancer phenotype; the targeted detection and quantification of mutated proteins can help reveal the mechanisms of mutation-driven cancers and could be used for cancer diagnosis.^16^ The current state of the art of targeted mass spectrometry of mutated proteins is time intensive, requiring the purchase of synthetic standard peptides, PRM assay development, and sequential running of unlabeled peptide samples.^16^ With GoDig-LiF, we envisioned a workflow that skips most of these steps, requiring only the sample, mass spectrometer, and knowledge of the mutations, and that quantifies targets in several samples in a single run (Fig. 2B).

**Figure 2.**
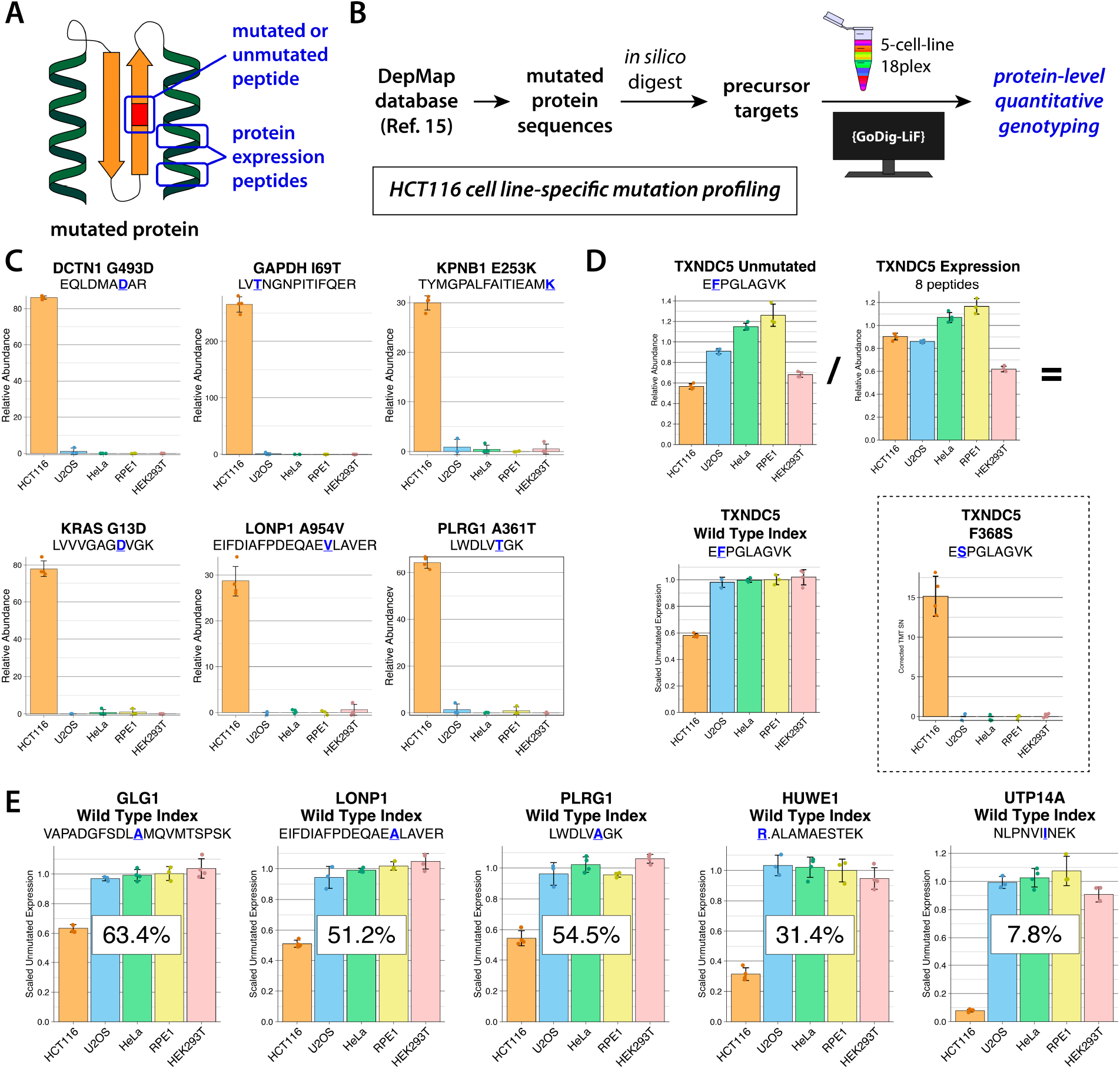
Quantitative protein-level genotyping using GoDig-LiF. **A.** Diagram of mutated, unmutated, and protein expression peptides on a hypothetical protein. **B**. GoDig-LiF workflow for quantitative protein-level genotyping. **C**. Examples of quantification of mutated peptides in a 5-cell-line 18plex. **D**. Illustration of expression peptide normalization to remove the effects of cell line-specific whole protein expression levels. **E**. Examples of wild type index quantification.

We accessed the DepMap portal^17^ to retrieve single-amino acid mutations in the HCT116 cell line and performed GoDig-LiF, targeting and quantifying 26 mutated peptides in a previously described 5-cell-line 18plex sample (Fig. 2B–C, Fig. S7).^8^ All 26 of these peptides exhibited the expected quantitative pattern, showing >5x intensity in the HCT116 channels compared to the others. These 26 assays are validated, publicly available, and portable, and were developed with an unprecedentedly low level of prior investment. Notably, we were able to quantify KRAS G13D, a known driver of cancer.^16,18^

Whereas mutated peptides indicate the presence of a mutated protein, we sought to quantify the penetration of these mutations to the protein level by adding to our assays the unmutated versions of the mutated peptides and “expression peptides” that do not cover the mutation site, measuring the expression level of the protein (Fig. 2A). Normalizing by protein expression allows an estimation of the proportion of the proteins that are unmutated; we call this estimate the “wild type index” (Fig. 2D). From our 26 validated mutations, 14 with peptides specific to the mutated and unmutated forms were selected; for each of these, we targeted both, as well as expression peptides passing strict *in silico* digest filters, limited to the 10 most detectable per protein according to our random forest-based model (Fig. 2D, Table S5). We quantified 9 wild type indices, and all 9 showed the expected quantitative pattern: a reduced amount of unmutated protein in HCT116 compared to the other cell lines (Fig. 2E, Fig. S8, Table S6). Unlike with DNA-level genotyping, this assay is able to quantify mutation at the protein level; for example, with the homozygous UTP14A I669V mutation, we measured a wild type index of 7.8%, showing good agreement with the 96.3% allele frequency reported by DepMap.^17^

In conclusion, we used the Prosit-TMT model to enable targeted multiplexed proteomics with nothing but a TMTpro-labeled sample, a target list, the freely available GoDig-LiF software, and an Orbitrap Eclipse or Ascend mass spectrometer. We used a new Comet rank filter to reduce the identification FDR to ∼0.1%, established empirical filters for making *in silico* digests efficient, trained a detectability prediction model for peptide selection, and validated our approach by genotyping samples at the protein level with unprecedented ease. GoDig-LiF can be used to target any protein or peptide of interest, regardless of prior observations, while leveraging the sensitivity and high sample throughput of TMT-based targeted mass spectrometry.

## METHODS

### Sample Preparation

Only aliquots of previously published samples were used for this study: the 5-cell-line mixture and the human- and yeast-derived samples for the FDR estimation were described in Shuken et al., *Anal. Chem*. **2023**, *95*, 15180. Lyophilized peptides were resuspended in 3% acetonitrile + 1% formic acid in water and 500 ng of peptide was injected per run.

### In Silico *Digest and Predicted Library Generation*

*In silico* digests were performed in R using the stringr package. Generating a predicted library in GoDig-LiF requires a set of target peptide predictions and a set of background peptide predictions; all predicted libraries used in this study used the same background predictions, i.e., a strict *in silico* digest of the human proteome. For this broadly applicable human *in silico* digest, peptide length was in [7, 25], no missed cleavages were allowed, no Met oxidations were allowed, only z = 2 precursors were included, m/z (Th) was in [500, 1200], and only peptides with charge = 2 in acidic buffer (1 + # His + # Lys + # Arg) were included. Only the canonical human reference proteome from Uniprot (UP000005640_9606.fasta without “additional.fasta” file) was included.

For the 18 HSP90A peptides, 18 precursors were selected based on being identified in all of 4 untargeted DDA runs of a human sample (Fig. 1C, Table S1).

For the 97 HSP90A precursors, HS90A_HUMAN (Uniprot P07900) was digested with the following parameters: peptide length was in [7, 30], missed cleavages were in [0, 1], Met oxidations were in [0, 1], z was in [2, 4], m/z (Th) was in [400, 1400], and charge in acidic buffer was in [1, 3].

For the permissive *in silico* digest used in Fig. 1D, the full human reference proteome from Uniprot (UP000005640_9606.fasta + UP000005640_9606_additional.fasta) was digested with the following parameters: peptide length was in [7, 30], missed cleavages were in [0, 2], Met oxidations were in [0, 3], z was in [2, 5], m/z (Th) was in [400, 1400], and charge in acidic buffer was in [1, 6]. We assembled this *in silico* digest into metapeptides (Fig. S6). We then compiled all precursors identified in the two experimental datasets described in Fig. 1D (Shuken et al., *J. Proteome Res*. **2024**, *23*, 1834) into a single experimental list and then discarded all metapeptides that did not contain precursors in that experimental list. The resulting precursors, all generated by permissive *in silico* digest and existing in metapeptides containing experimentally identified precursors, were used for the comparisons in Fig. 1D. For the *in silico* digest used to train the detectability model (Fig. 1E), see below.

For the *in silico* digest of mutated protein sequences from DepMap, see below.

### Detectability Model

The peptide detectability model was trained using the PeptideRanger package in R (Riley et al., *J. Proteome Res*. **2023**, *22*, 526). To train the model on peptides that are difficult to distinguish using filters, we used the stringent *in silico* digest parameters illustrated in Fig. 1E to generate a set of likely detectable peptides from the full human reference proteome downloaded from Uniprot (UP000005640_9606.fasta + UP000005640_9606_additional.fasta): peptide length was in [7, 25], no missed cleavages were allowed, no Met oxidations were allowed, z was in [2, 3], m/z (Th) was in [450, 1100], charge in acidic buffer was in [1, 3], and mobile proton index was in [0, 1]. As described above, we assembled this *in silico* digest into metapeptides and then discarded all metapeptides not containing experimentally identified precursors. For training, the ‘n_obs_pep’ and ‘ratio’ parameters were set to 1 for experimentally identified precursors and 0 for unidentified precursors, and the ‘n_obs_prot’ parameter was set to 1 for all precursors. The ROC plot in Fig. 1E is of a randomly selected 25% of the data predicted with a model trained on the other 75%, but a model trained on 100% of the data was used to select expression peptides for wild type index measurements.

### *Retrieval*, In Silico *Digest, and Selection of Mutated and Unmutated Peptides*

All HCT116 mutations were downloaded from 23Q4 DepMap database (OmicsSomaticMutations.csv) at https://depmap.org/portal/data_page/?tab=allData. Missense variants were retained and all other mutations were discarded. **L→I and I→L** mutations were removed. Only entries with Uniprot IDs were retained and the protein sequences were retrieved using the UniProt.ws R package. Incorrect entries whose protein sequences did not have the indicated amino acid at the indicated position were discarded and then the mutation was applied to the protein sequence *in silico*. Each form of each protein was then digested *in silico* with the following parameters: length was in [7, 25], no missed cleavages were allowed, no Met oxidations were allowed, z was in [2, 3], m/z (Th) was in [450, 1100], charge in acidic buffer was in [1, 3], and mobile proton index was in [0, 1]. Only peptides matching one form of the protein and not the other were retained. The peptides were then cross-referenced to the permissive human *in silico* digest described above (used in Fig. 1D), and peptides with any gene-level ambiguity were discarded. The remaining precursors were targeted with GoDig-LiF as described below and in the main text.

### Liquid Chromatography-Tandem Mass Spectrometry

All LC-MS/MS experiments were performed on an Orbitrap Eclipse mass spectrometer (Thermo Fisher Scientific) equipped with an EASY-nLC 1200 system (Thermo Fisher Scientific) bearing a 30-cm column packed with ReproSil-Pur C18-coated beads that are 2.4 µm in diameter (Dr. Maisch GmbH, part no. r124.aq.0001) according to the FlashPack protocol (Kovalchuk, Jensen, and Rogowska-Wrzesinska, *Mol. Cell. Proteomics* **2019**, *18*, 383) and heated to 60 °C. The column was mounted onto a nanospray ion source designed and constructed in-house and operated in positive mode at 2600 V with the transfer tube heated to 300 °C. All spectra were acquired in centroid mode with the RF Lens set to 30% amplitude.

For the HSP90A analyses, 0.5 µg of peptide was injected and separated using the following 1-h gradient at 500 nl/min: a linear increase from 2% Buffer B (5% water + 0.125% formic acid in acetonitrile) in Buffer A (5% acetonitrile + 0.125% formic acid in water) to 15% Buffer B over 39 min, followed by a linear increase to 27% Buffer B over 13 min, followed by a linear increase to 100% Buffer B over 4 min, followed by isocratic flow at 100% Buffer B for 4 min.

For all other analyses, 0.5 µg of peptide was injected and separated using the following 2-h gradient at 500 nl/min: a linear increase from 5% Buffer B (5% water + 0.125% formic acid in acetonitrile) in Buffer A (5% acetonitrile + 0.125% formic acid in water) to 28% Buffer B over 97 min, followed by a linear increase to 44% Buffer B over 13 min, followed by a linear increase to 100% Buffer B over 5 min, followed by isocratic flow at 100% Buffer B for 5 min.

The DDA library-based HSP90A GoDig analyses were performed with the following parameters. The published 24-fraction 4-cell-line based library was used (Shuken et al., *J. Proteome Res*. **2024**, *23*, 1834). A bin width of 0.5 min was used to establish 240 bins from the 120-min library data and a monitor window of ±10 bins was used so that during the 60-min GoDig run, on average, target monitoring began 2.5 min before elution and ended 2.5 min afterwards (60 min / 240 bins = 0.25 min/bin). A monitor dynamic exclusion duration of 5 s was used. Ion trap MS2 (ITMS2) monitoring was performed with an isolation width of 0.5 Th; a maximum injection time (max IT) of 120 ms and automatic gain control (AGC) target of 10^4^ was used; ion trap CID was performed with a normalized collision energy (NCE) of 35%; the Normal scan rate setting was used; and a fragment ion matching tolerance of 0.15 Th was used. For IDMS2 scans, CID was performed with an NCE of 35.1%; an AGC target of 10^5^ was used; a max IT of 900 ms was used; the orbitrap was operated at Resolution = 15k; the fragment match tolerance was 15 ppm; and an MS3 was queued only if ≥4 fragment peaks were matched AND the cosine similarity score exceeded 0.9. Ion trap MS3 prescans were enabled. For orbitrap MS3 scans, a max IT of 1,000 ms was used; fragment ions 50 Th below and 5 Th above the precursor m/z were excluded from SPS; 4 SPS ions were isolated for MS3 using an isolation width of 0.8 Th; HCD was performed with an NCE of 65; and the orbitrap was operated at Resolution = 50k. Dynamic close-out was disabled.

Unless otherwise specified, GoDig-LiF analyses were performed with the same parameters as those described above for the DDA library-based HSP90A analysis, except that a bin width of 0.75 min was used to establish ∼240 bins from the predictions which span ∼180 min of iRT values. For mutation experiments, 6 priming runs (Shuken et al., *J. Proteome Res*. **2024**, *23*, 1834) were performed before the analytical run.

### Data Processing and Analysis

Unless otherwise specified, GoDig data were processed in the following manner. MS3-level data were extracted from the raw files using the Data Analysis tab of the GoDig software. In R, MS3-level data were then collapsed to the precursor, mutation site/form, or protein level by summing the signal-to-noise ratios in each channel. An analyte (precursor, peptide, or site) was considered quantified only if the SumSN exceeded 10 * (# channels); for quantitative analysis, any analyte that failed to exceed this was discarded. In R, quantitative data were then corrected using the isotopic purity values contained in the certificate of analysis for the TMT reagent lot, publicly available on www.thermofisher.com.

Comet searches were performed either *post hoc* using an in-house pipeline (for direct comparisons) or were integrated into the GoDig-LiF software using the publicly available Comet source code (https://github.com/UWPR/Comet); in either case, the following search parameters were used. Cysteine carbamidomethylation and TMT labeling of peptide N-termini and K were set as static modifications and M oxidation was used as a variable modification. The precursor m/z tolerance was 50 ppm. The fragment m/z tolerance was 0.02 Th.

### Software Development

GoDig-LiF feature implementation and debugging was done in C# using Visual Studio 2022 (Microsoft).

## DATA AVAILABILITY

All raw data, GDVXML files, search results, *in silico* digests, Prosit-TMT predictions, GoDig outputs, and GoDig libraries (data-or prediction-based) generated or used in this work have been deposited in the PRIDE (Perez-Riverol, et al., *Nucleic Acids Res* **2022**, *50*, D543) repository via the ProteomeXchange Consortium with the identifier PXD066607.

## CODE AVAILABILITY

The GoDig-LiF software is freely available via a Recipient Agreement for the Orbitrap Eclipse or Orbitrap Ascend mass spectrometer with an iAPI license. The Recipient Agreement and instructions for obtaining the iAPI license can both be obtained at https://gygi.hms.harvard.edu/software.html. GoDig-LiF is enabled through the iAPI framework such that the GoDig-LiF source code is not available, but iAPI sample code is available at https://github.com/thermofisherlsms/iapi. GoDig uses Comet (ver. 2025.02.0), available at https://uwpr.github.io/Comet.

## ACKNOWLEDGMENTS

The authors thank members of the Gygi Laboratory for helpful conversations and support. This work was funded in part by the National Institutes of Health (NIH) grants AG088297 (S.R.S.) and GM67945 (S.P.G.).

## AUTHOR CONTRIBUTIONS

S.R.S. and S.P.G. conceived the project. S.R.S. wrote all code, prepared all samples, performed all LC-MS/MS experiments, processed and analyzed all data, made all figures, and wrote the manuscript. S.R.S. and S.P.G. edited the manuscript.

## Notes

https://www.ebi.ac.uk/pride/archive/projects/PXD066607

### Competing Interest Statement

The authors have declared no competing interest.

## REFERENCES

1 Marx, V. “Targeted proteomics.” Nature Methods 2013, 10, 19–22.

2 Antelo-Varela, M.; Bumann, D.; Schmidt, A. “Optimizing SureQuant for Targeted Peptide Quantification: a Technical Comparison with PRM and SWATH-MS Methods.” Anal. Chem. 2024, 96, 18061–18069.

3 Yang, K.; Paulo, J. A.; Gygi, S. P.; Yu, Q. “Enhanced Sample Multiplexing-Based Targeted Proteomics with Intelligent Data Acquisition.” J. Am. Soc. Mass Spectrom. 2024, 35, 2420–2428.

4 Remes, P. M.; Yip, P.; MacCoss, M. J. “Highly Multiplex Targeted Proteomics Enabled by Real-Time Chromatographic Alignment.” Anal. Chem. 2020, 92, 11809–11817.

5 Zhu, H.; et al. “PRM-LIVE with Trapped Ion Mobility Spectrometry and Its Application in Selectivity Profiling of Kinase Inhibitors.” Anal. Chem. 2021, 93, 13791–13799.

6 Yu, Q.; et al. “Sample multiplexing-based targeted pathway proteomics with real-time analytics reveals the impact of genetic variation on protein expression.” Nature Comm. 2023, 14, 555.

7 Dong, K. D.; Schmid, E. W.; Bomgarden, R. D.; Choi, J. H.; Gygi, S. P.; Yu, Q.; Paulo, J. A. “Adapting an Isobaric Tag-Labeled Yeast Peptide Standard to Develop Targeted Proteomics Assays.” J. Proteome Res. 2024, 23, 142–148.

8 Shuken, S. R.; Yu, Q.; Gygi, S. P. “Inserting Pre-analytical Chromatographic Priming Runs Significantly Improves Targeted Pathway Proteomics with Sample Multiplexing.” J. Proteome Res. 2024, 23, 1834–1843.

9 Gabriel, W.; Giurcoiu, V.; Lautenbacher, L.; Wilhelm, M. “Preedicting fragment intensities and retention time of iTRAQ- and TMTPro-labeled peptides with Prosit-TMT.” Proteomics 2022, 22, 2100257.

10 Shuken, S. R. “An Introduction to Mass Spectrometry-Based Proteomics.” J. Proteome Res. 2023, 22, 2151–2171.

11 Eng, J. K.; Jahan, T. A.; Hoopmann, M. R. “Comet: An open-source MS/MS sequence database search tool.” Proteomics 2013, 13, 22–24.

12 Zhang, B.; Chambers, M. C.; Tabb, D. L. “Proteomic Parsimony through Bipartite Graph Analysis Improves Accuracy and Transparency.” J. Proteome Res. 2007, 6, 3549–3557.

13 Shuken, S. R.; et al. “Limited proteolysis-mass spectrometry reveals aging-associated changes in cerebrospinal fluid protein abundances and structures.” Nature Aging 2022, 2, 379–388.

14 Paizs, B.; Suhai, S. “Fragmentation pathways of protonated peptides.” Mass Spectrom. Rev. 2005, 24, 508–548.

15 Riley, R. M.; et al. “PeptideRanger: An R Package to Optimize Synthetic Peptide Selection for Mass Spectrometry Applications.” J. Proteome Res. 2023, 22, 526–531.

16 Lin, T.-T.; et al. “Mass spectrometry-based targeted proteomics for analysis of protein mutations.” Mass Spec. Rev. 2023, 42, 796–821.

17 DepMap, Broad. “DepMap 23Q4 Public Figshare+ Dataset.” 2023, 10.25452/figshare.plus.24667905.v2.

18 Matsunaga, K.; et al. “Clinical significance of the KRAS G13D mutation in anastomotic recurrence of colorectal cancer.” Oncol. Lett. 2023, 25, 192.

