## Supporting Information for "Library-Free Multiplexed Targeted Proteomics Enables Quantitative Protein-Level Genotyping"

**SUPPLEMENTARY FIGURES S1–S8 .......... 2**

### Supplementary Figures S1–S8

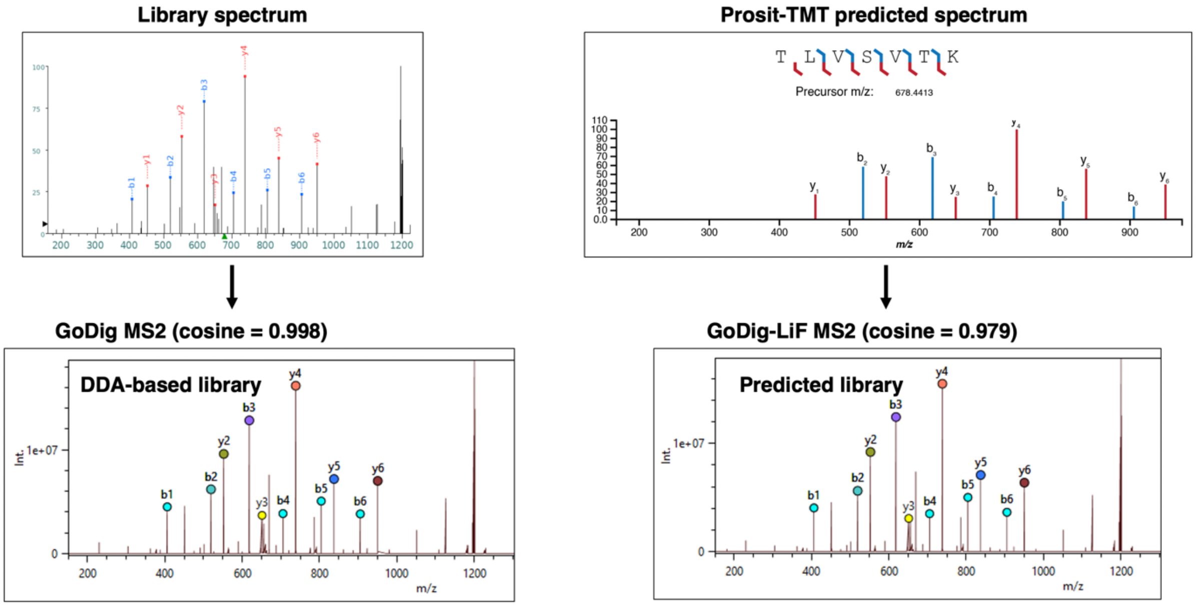


**Figure S1.** Examples of library, predicted, and empirical MS2 spectra acquired during GoDig and GoDig-LiF analysis (experiment depicted in Figure 1B).

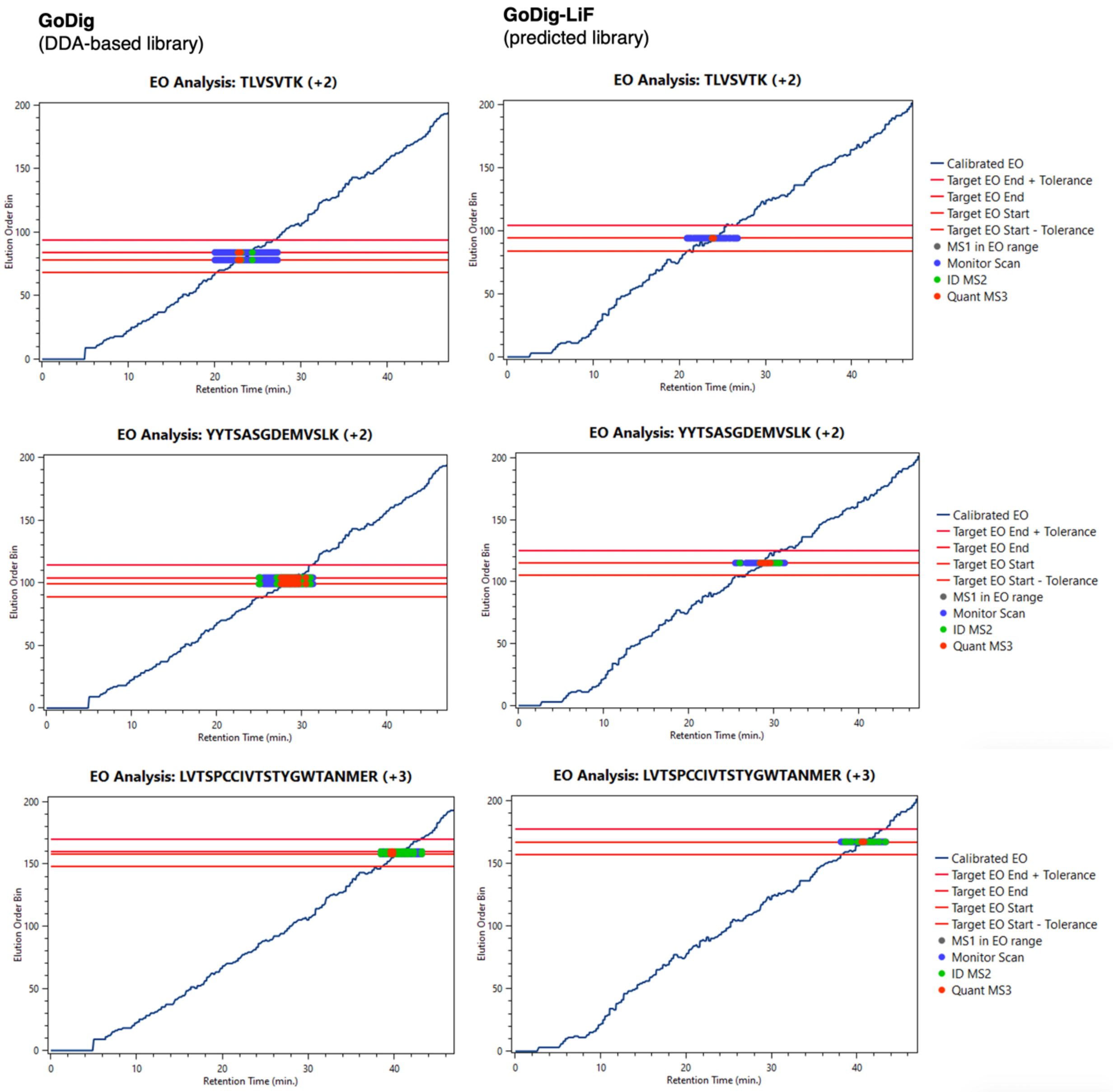


**Figure S2.** Examples of EO prediction with GoDig-LiF compared to the DDA-based GoDig library (experiment depicted in Figure 1B).

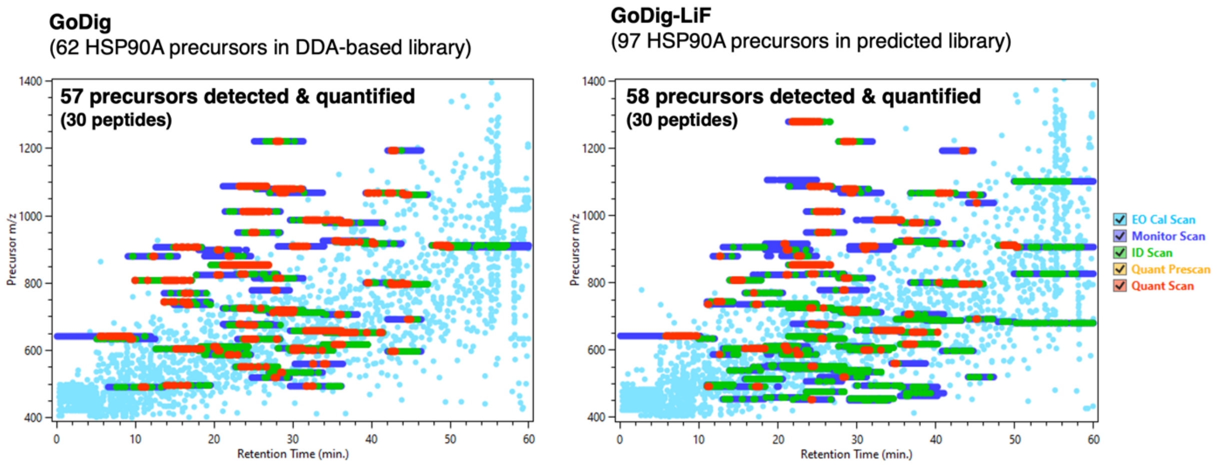


**Figure S3.** GoDigViewer visualization of GoDig and GoDig-LiF analysis of HSP90A.

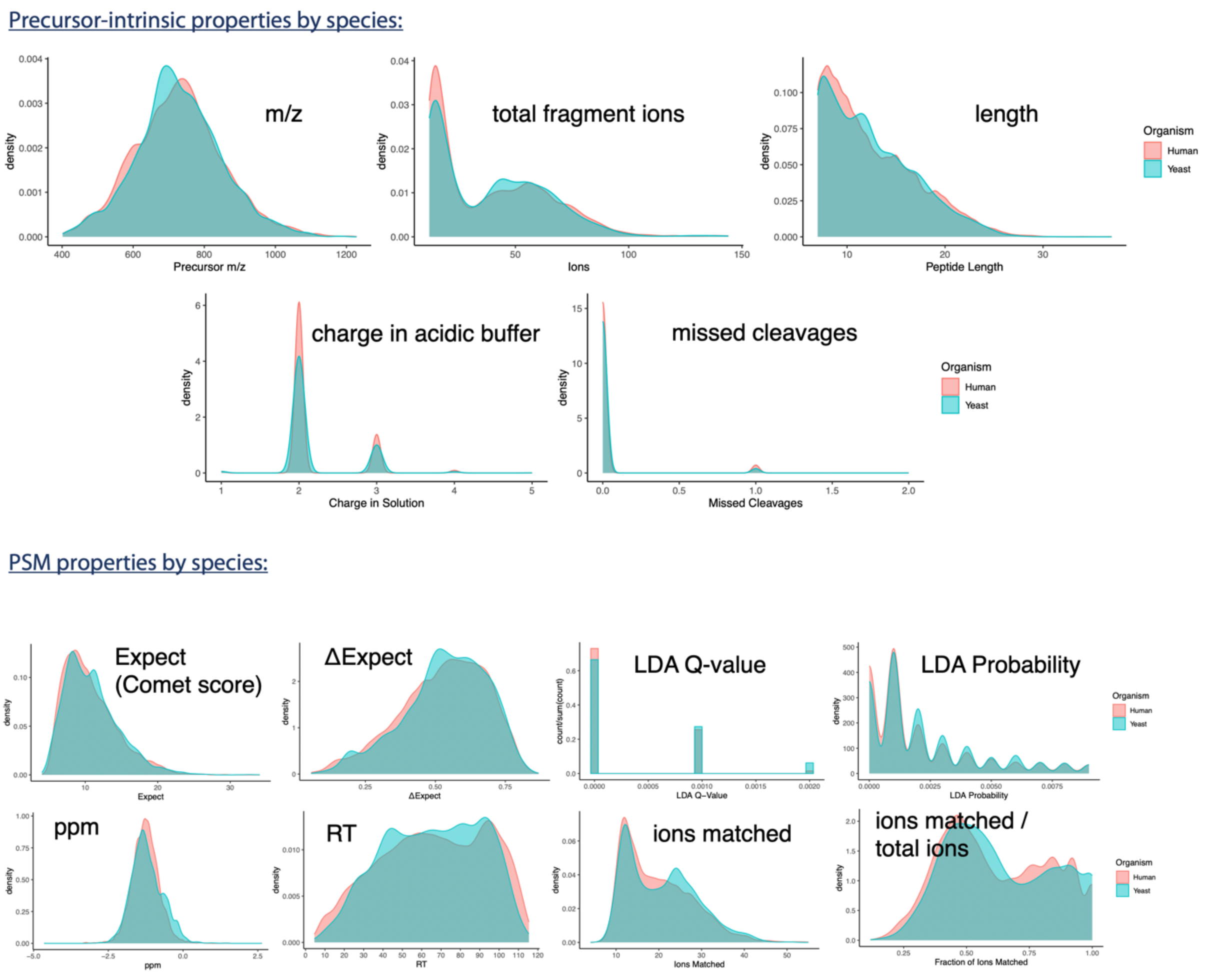


**Figure S4.** Properties of 7,008 human-specific and 3,982 yeast-specific precursors as well as the PSMs in which they were identified. LDA = linear discriminator analysis.

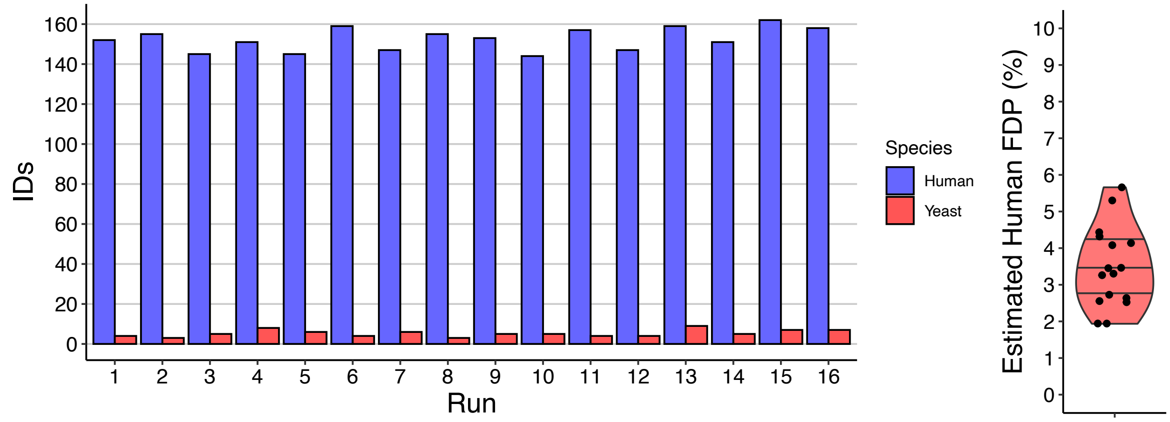


**Figure S5.** Estimation of the false discovery rate (FDR) of GoDig-LiF without the Comet rank filter.

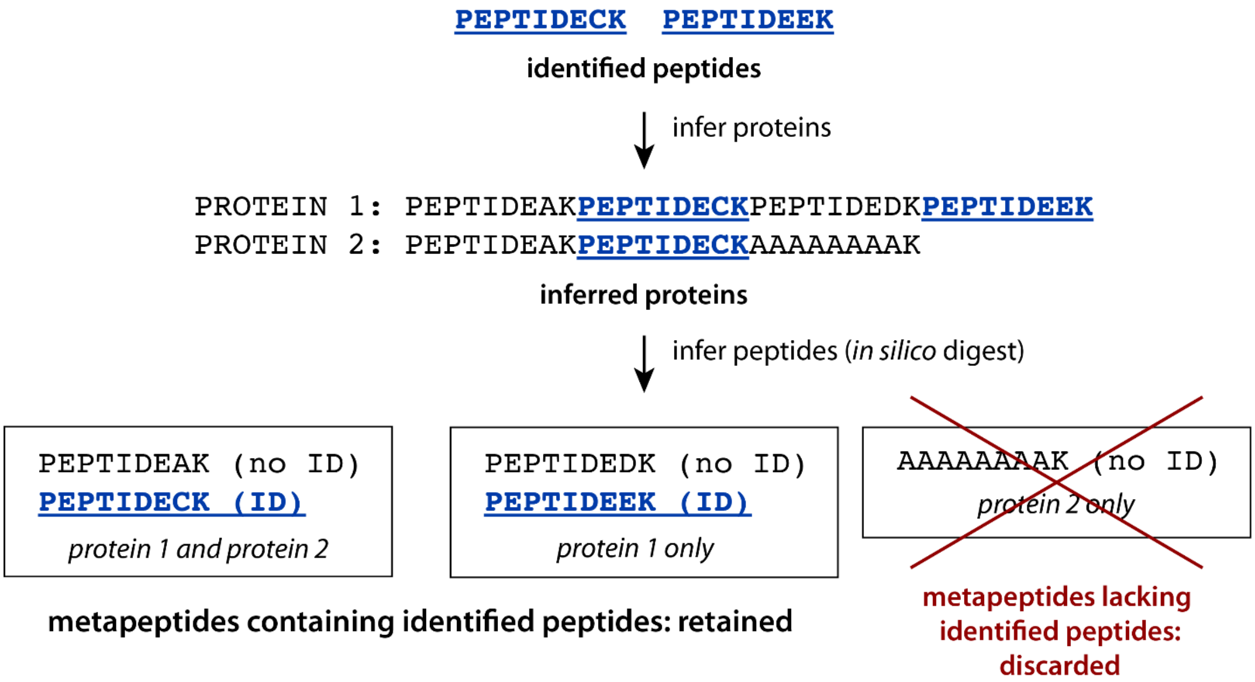


**Figure S6.** Procedure for generating identified and unidentified precursors as training data for the detectability model. The metapeptide protein (and peptide) grouping method is described in Shuken et al., *Nature Aging* **2022**, *2*, 379, based on the metapeptide term coined in Zhang, Chambers, and Tabb, *J. Proteome Res.* **2007**, *6*, 3549. The metapeptide method results in mutually exclusive sets of peptides where all peptides in a given set match the same set of proteins. As a result of this procedure, the existence in the sample of each unidentified peptide is implied by the fact that at least one peptide in the same set (metapeptide) was identified. In this example, the observed peptides do not prove that protein 2 existed in the sample, so AAAAAAAAK, which matches only protein 2, is discarded from the *in silico* digest.

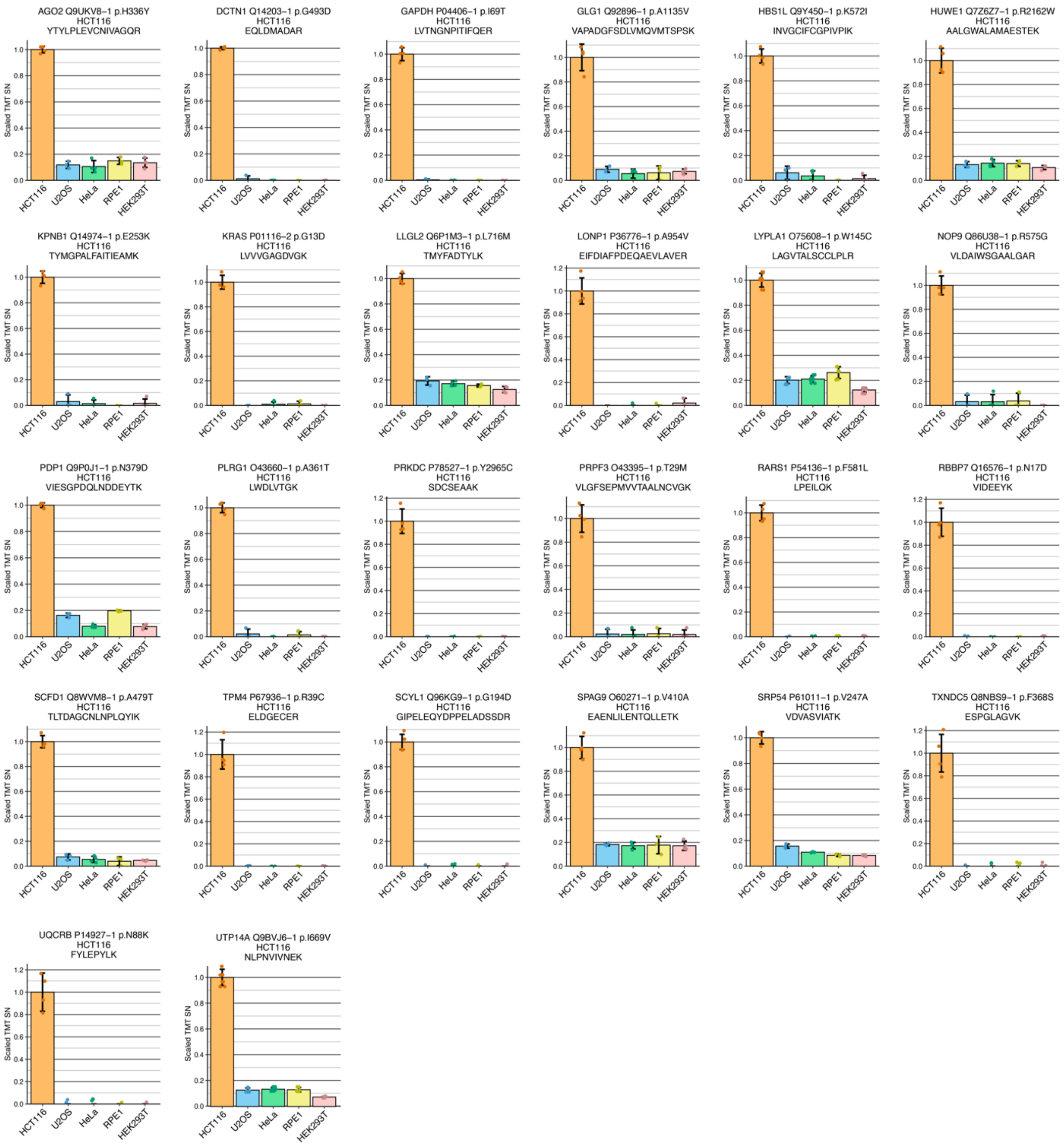


**Figure S7.** Quantification of all 26 mutated peptides.

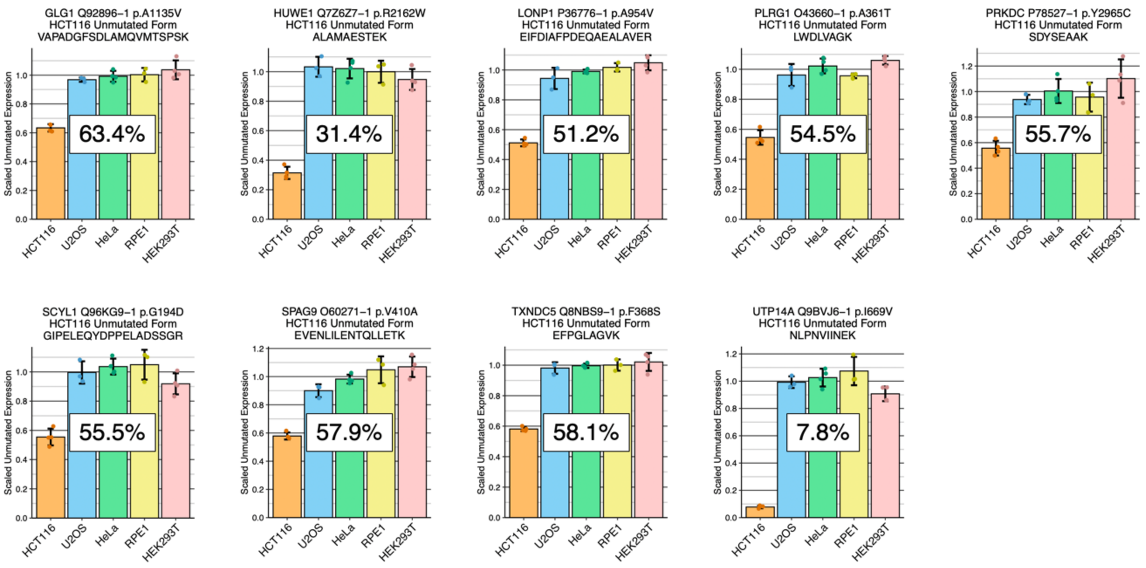


**Figure S8.** Quantification of all 9 wild type indices. The wild-type index reports the fraction of a protein’s population that is the wild-type form of a protein compared to the mutated form.
